# Signalling maps in cancer research: construction and data analysis

**DOI:** 10.1101/089409

**Authors:** Maria Kondratova, Nicolas Sompairac, Emmanuel Barillot, Andrei Zinovyev, Inna Kuperstein

**Affiliations:** Institut Curie, 26 rue d’Ulm, F-75005 Paris, France, Inserm, U900, F-75005, Paris France, Mines Paris Tech, F-77305 cedex Fontainebleau, France, PSL Research University, F-75005 Paris, France

**Keywords:** Knowledge formalisation, cancer signalling map construction, data model, map navigation, data analysis and visualization, biocuration, graphical standard

## Abstract

Generation and usage of high-quality molecular signalling network maps can be augmented by standardising notations, establishing curation workflows and application of computational biology methods to exploit the knowledge contained in the maps. In this manuscript, we summarize the major aims and challenges of assembling information in the form of comprehensive maps of molecular interactions. Mainly, we share our experience gained while creating the Atlas of Cancer Signalling Network. In the step-by-step procedure, we describe the map construction process and suggest solutions for map complexity management by introducing a hierarchical modular map structure. In addition, we describe the NaviCell platform, a computational technology using Google Maps API to explore comprehensive molecular maps similar to geographical maps, and explain the advantages of semantic zooming principles for map navigation. We also provide the outline to prepare signalling network maps for navigation using the NaviCell platform. Finally, several examples of cancer high-throughput data analysis and visualization in the context of comprehensive signalling maps are presented.

## Introduction

Similar to geographical maps, the representation of biological knowledge as a diagram facilitates the study of complex processes in the living cell, in a visual and insightful way.

### Goal of knowledge formalisation

The representation of biological processes as comprehensive signalling network maps has three major goals: (i) to generate a resource containing a formalised summary of biological findings from many research groups, (ii) to provide a platform for sharing information and discussing biological mechanisms, (iii) to create an analytical tool useful for high-throughput data integration and analysis.

It can be helpful to systematically represent and formalise the molecular information distributed in thousands of scientific publications. An additional advantage of representing biological processes in a graphical form is to capture the multiple cross-talks and interactions occurring between different cell processes (1).

Analysis and visualisation of omics data in the context of signalling network maps can help to detect patterns in the data projected onto the molecular mechanisms there represented. For instance, identification of deregulated mechanisms and key players in human diseases, has a direct clinical application (2)(3). Moreover, correlating the status of those deregulated mechanisms with patient survival, helps for patient stratification according to their network-based signatures (4). Due to the complexity of mechanisms simultaneously involved in diseases, targeting combinations of molecular players is now the trend in treatment of complex diseases. The computational approaches using signalling maps allow testing multiple combinations *in silico*, considering large comprehensive signalling networks and omics data(5)(6). In addition, signalling networks can serve for modelling and prediction of cell fate decisions, (7)(8) and suggestion of non-intuitive combinations of gene perturbations to explain phenotypes in health and disease (9). To achieve these goals, the construction of a signalling map should become an accessible procedure that can be completed in a reasonable time. There are several solutions for biological knowledge formalisation briefly described in this manuscript. We contribute to this global aim and formulate the main principles and steps of the established workflow for manual map construction. In addition, we suggest the biological network map navigation facilitated by Google Maps technology and provide examples of data analysis and visualisation in this context.

### Diagram types for molecular processes representation

Generally speaking, there are four main approaches (or diagram types) for representing molecular processes, each of them characterized by a certain depth of description: (i) *interaction diagram*, which shows simple binary relations between molecular entities; (ii) *activity-flow*, known as regulatory network or influence diagrams, representing the flow of information or influences of one entity on another; (iii) *entity relationship diagram*, depicting relations in which a given entity participates, and (iv) *process description (PD) diagram*, known in chemical kinetics as bi-partite reaction network graphs (10).

### Pathways and network maps approached for molecular processes representation

Using the aforementioned approaches of molecular processes representation, several pathway databases have emerged (11). They serve as biological knowledge information resource and as computational analytical tools for systems-based interpretation of data. A significant number of pathway databases has been also developed in the private domain, but the majority of pathway collections are free and open source (Supplementary Table 1).

The most common way of knowledge representation, used in the majority of these resources, is depicting separate processes referring to ‘signal transduction pathways’. However, drawing individual pathways precludes clear representation of cross-regulations among biological processes. The alternative solution is to create seamless maps of biological mechanisms covering multiple cell processes simultaneously, similar to geographical maps. In order to apply this “geography-inspired” approach to biological knowledge, it is necessary to address a number of challenges related to generation, maintenance and navigation of large signalling network maps. We discuss these challenges and suggest solutions based on our long-term experience manipulating biological maps.

### Common standards for molecular processes representation

With the aim of creating a collection of exchangeable comprehensive signalling maps, common rules for drawing maps and standard graphical syntax should be developed and consistently applied. The current suggested solution in the field is the Systems Biology Graphical Notation (SBGN) syntax. This syntax is compatible with various pathway drawing and analytical tools, allowing to represent not only biochemical processes, but also cell compartments and phenotypes (10). Furthermore, to increase cross-compatibility between pathway resources and analytical tools, several common formats for exchanging information on molecular interactions, such as BioPAX, SBML, PSI-MI etc., have been suggested (7).

### Tools for molecular processes representation

Since the generation of signalling maps is a long and laborious process, the choice of the appropriate drawing tool, best suited for the type of network, has to be thoughtfully done. There are several free and commercial tools to create biological network diagrams that differ in the process representation approach, syntax and requirement for the end users’ technical skills, as SBGN-ED (12), visANT (13), CellDesigner (14), etc. (Supplementary Table 2).

### Navigation platforms for comprehensive molecular network maps

Visualising and exploring biological network diagrams became an important issue, because size and complexity of molecular networks reach the dimensions of modern geographic maps. Therefore, several tools such as CellPublisher (15), Pathways projector (16) MINERVA (17), NaviCell (18)(19) and others, have adopted the navigation logic from Google Maps technology. These tools allow exploring big networks in a user-friendly manner thanks to such Google Maps features, as scrolling, zooming, markers and callouts (Supplementary Table 2).

### A workflow for construction of comprehensive signalling network maps

In this manuscript we describe a set of good practices for building comprehensive signalling network maps. We provide the methodology that allows to overcome challenges associated with construction, navigation and exploration of large molecular interaction maps (Figure 1). We suggest an approach that is neither unique nor universal, but provides verified practical solutions to comprehensive map construction and manipulation that successfully served to generate the ACSN resource (20), and also applied in other studies (21)(9).

**Figure 1.**
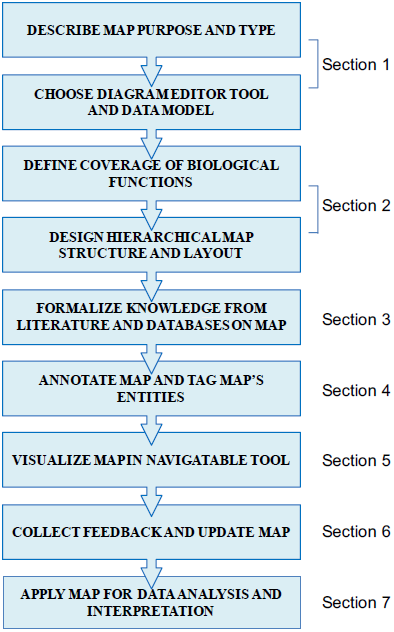
Map construction workflow scheme.

The corresponding section in the texts is indicated.

The suggested workflow is in line with the FAIR principles for data management and sharing (23).The access to the detailed procedures is provided in the Supplementary materials and available at https://github.com/sysbio-curie/NaviCell. The terms and definitions used through the paper are listed in Table 1.

**Table 1.** **terms and definitions.**

| Graphical standards and exchange formats terms and definitions |  |
| --- | --- |
| SBGN | Systems Biology Graphical Notation (SBGN) is a standard graphical syntax for representation of biological processes and interactions. SBGN is compatible with multiple pathway drawing and analytical tools, http://sbgn.github.io/sbgn/ |
| SBML | Systems Biology Markup Language (SBML) is a representation format, based on XML, for communicating and storing computational models of biological processes. It is a free and open standard language with widespread software support, http://sbml.org |
| Standard identifier (ID) | Community-accepted nomenclature for scientific naming of biomolecules as genes, proteins, chemicals, drugs etc. The sources for standard IDs are repositories as UNIPROT, CHEB, HUGO, http://identifiers.org |
| Data and models exchange formats | Standard formats for data and models to facilitate networks and software interoperability. There are two major standard networks exchange formats, BIOPAX for complex networks and SIF for simple binary interactions. The CellDesigner xml format is commonly-used exchange format compatible with multiple network analysis tools. |
| Signalling network map terms and definitions |  |
| Map (in ACSN) | Diagram of detailed molecular interactions with meaningful layout reflecting a certain biological process, which is graphically represented in CellDesigner tool and converted to NaviCell format for exploration and curation. |
| Map layer (in ACSN) | Map area covering several molecular processes with similar functions. |
| Map module (in ACSN) | Part of the map representing a sequence of molecular interactions responsible for execution of a particular function. |
| Signal transduction pathway or signalling cascade | Sequence of molecular interaction that transforms extracellular signals into an intercellular activity. |
| Map entity (in ACSN) | Component of the map graphically depicted using SBGN standards in CellDesigner tool. |
| Hierarchical structure of a map | System for signaling network map complexity reduction by dividing it into **hierarchically organized units** as layers meta-modules, modules that exist as independent sub-maps. The structure supports the **horizontal map navigation mode**. |
| Confidence score (ACSN) | Value representing the measure of accuracy of binary interactions on the map. There are two confidence scores in NaviCell maps. The **reference score** indicates number and “weight” associated to the publications found in the annotation of a given reaction. The **functional proximity score** is computed based on the external network of protein-protein interactions (PPI), Human Protein Reference Database (HPRD). |
| NaviCell terms and definitions |  |
| Semantic zoom | A mechanism providing several map views with different levels of details depiction achieved by gradual exclusion of details while zooming out. It simplifies navigation through large maps of molecular interactions by providing several levels of details, resembling navigation through geographical maps. Exploring the map from a detailed toward a top-level view is achieved by gradual exclusion and modification (simplification and abstraction) of details. One of the main principle of semantic zooming is in that every detail which is shown on the map at a current zoom level, should be readable. This feature supports the **vertical map navigation mode**. |
| Bird-eye view panel | Window containing top-level view of the map with indication on currently centered area; adapted from Google maps. |
| Zooming bar | Zoom control slider; adapted from Google maps. |
| Marker | Symbol indicating location of chosen objects on the map; adapted from Google maps. |
| Pop-up bubble | Small window that opens by clicking on marker. Contains short description and hyperlinks related to the marked entity. |
| Annotation post | Detailed map entity annotation created in CellDesigner by map manager. The annotation is converted to annotation post and displayed in the associated blog by NaviCell. |

The DNA repair map from ACSN resource is used as an example for map construction and annotation: https://acsn.curie.fr/navicell/maps/dnarepair/master/index.html (20).

EMT regulation map from NaviCell collection (9) is used for data visualisation: https://navicell.curie.fr/pages/signalling_network_emt_regulation_description.html

## Section 1: General principles for map construction

### 1.1 Work organization

Signalling map construction requires an overview of broad scientific literature and a very meticulous work for correct representation of molecular processes in great detail. Therefore, several important decisions should be made prior to the map construction: (i) What is the purpose for constructing the map? (ii) What diagram type is suitable to properly represent the knowledge? (iii) What is the appropriate tool to build the map? (iv) What processes should be included in the map? (iv) How the map will look like? Once those questions are answered, and an agreement on the approach has been reached, the signalling diagram construction should follow fixed principles to ensure generation of a homogeneous and accurate map. An additional important step before constructing a map is to consult similar efforts in the field, and evaluate the added value of a new map.

### 1.2 Map purpose and type

Referring to our example (DNA repair map), the purpose was to understand how different types of DNA damage are repaired in the cell and how these various DNA repair mechanisms and the cell cycle coordinate. To address this challenge and gather together these molecular mechanisms, we decided to construct a comprehensive map of DNA repair and cell cycle signalling. The map can be applied to **detect the modes of DNA repair machinery rewiring** in different pathological situations such as cancer or genotoxic stress.

The mechanisms of DNA repair are well studied and information on involved molecules and regulation circuits is available. Thus, to preserve and accurately depict the processes in its whole complexity, we have chosen the **process description (PD) diagram type,** where the biochemical reactions can be explicitly depicted.

### 1.3 Map construction tool, graphical standard and data model

Signalling processes of DNA repair are represented as biochemical reactions in CellDesigner diagram editor based on the **standard Systems Biology Graphical Notation (SBGN) syntax** (10), and the **Systems Biology Markup Language (SBML),** for further computational modelling of the map (14). The **data model** used in our example is schematically depicted in Figure 2. It includes such molecular entities as proteins, genes, RNAs, antisense RNAs, simple molecules, ions, drugs, phenotypes, and complexes. Edges on the map represent biochemical reactions or reaction regulations including post-translational modifications, translation, transcription, complex formation or dissociation, transport, degradation, etc. Reaction regulations are catalysis, inhibition, modulation, trigger, and physical stimulation. It is also possible to depict cell compartments such as cytosol, nucleus, mitochondria, etc. See http://celldesigner.org for Cell Designer tool guide and http://www.sbgn.org for SBGN syntax explanation.

**Figure 2.**
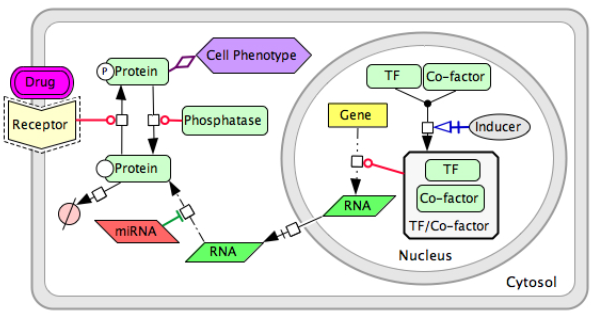
Data model for process description (PD) diagram using CellDesigner SBGN syntax.

## Section 2: Map boundaries, layout and structure

### 2.1 Map boundaries and content

Given the limitations of graphical tools and difficulties in manipulating large molecular interaction maps, the challenge is to define map boundaries. The solution that we suggest is to assign **one biological function per map** (e.g., cell cycle, angiogenesis, immune response). This is challenging per se due to “fuzziness” of borders between processes and overlaps between players and pathways. Therefore, biological function-driven maps can be assumed as components of a **global atlas** merged together via common players. To allow such combinable maps, common **standards for graphical representation** and **standard common identifiers** for naming the entities should be used through all maps. In addition, the community-based curation of maps is crucial for making more objective decisions regarding the map boundaries and content.

In our example, the boundaries and the content of the DNA repair map were defined in coordination with specialists in the corresponding fields. According to the seminal reviews and well-known databases, DNA repair machinery distinguishes 10 partially overlapping modes of repair, depending on the type of damage. The DNA damage types are depicted on the map as ‘inputs’ initiating the corresponding DNA repair mechanisms. In addition, the map covers four cell cycle phases and depicts regulatory circuits between cell cycle and DNA repair mechanisms coordinated by cell cycle checkpoints (24) (Figure 3A).

**Figure 3.**
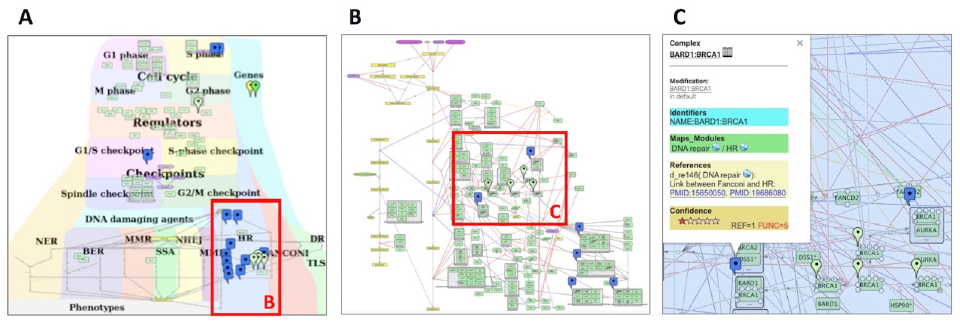
DNA repair map in NaviCell format. (A) Global layout of the map with maprker indicating BRCA1 protein distribution, (B) Individual module layout (Homologous recombination (HR), (C) Callout window with annotation of BARD:BRCA1 complex.

### 2.2 Map layout and hierarchical modular structure

The challenge of map dimensions and layout design should be addressed prior to the map drawing. This step is especially crucial in the case of collective map reconstruction where the final global map is assembled from a number of sub-maps generated by different map curators. The aim of signalling map construction is not only to summarize molecular mechanisms, but also to allocate the processes in a meaningful and biologically relevant way. Careful design of the **signalling map layout** helps for intuitive understanding of “what is where” and “what is close and what is distant”. In addition, a bird’s eye view on the map can give a general impression about the map complexity.

There are at least three options of map layout design: (i) Representing **spatial localization of processes** in the context of the global cell architecture. Most of the signal transduction maps resemble an accepted view of **cell organization**, they include cellular compartments and signalling pathways are placed in the corresponding compartments in the map (see examples at https://acsn.curie.fr). (ii) Depicting **process propagation in time**, for instance, demonstrating propagation of the signalling through the four phases of cell cycle (https://acsn.curie.fr/navicell/maps/cellcycle/master/index.html). (iii) Placing processes together according to their **involvement in a particular biological function** as DNA repair, where each pathway is depicting one biological function. These pathways are allocated next to each other, creating together a DNA repair machinery signalling (Figure 3A).

The combination of layout types at the same map is also possible as in the case of the DNA repair map, combining the three aforementioned layout types (Figure 3A and https://acsn.curie.fr/navicell/maps/dnarepair/master/index.html).

The global layout of the map can be also used as guidance to split it into sub-maps of layers and functional modules, and in generating a **hierarchical modular structure of the map** (see Table 1 for terms and definitions).

In our example, the **global map layout** of the DNA repair map has been designed to emphasize the **hierarchical organisation** of three major layers in the map: the DNA repair machinery layer and its connection to the cell cycle layer via the checkpoints layer. The upper layer depicts the cell cycle, the middle layer represents cell cycle checkpoints, receiving signals from the cell cycle and coordinating the crosstalk between the cell cycle and the DNA repair machinery, represented in the lower layer. This hierarchical organisation of the map in layers reflects the current understanding of signal propagation in the cell.

Further, the map can be divided into functional modules. Each functional module can exist as a part of the global map but also as an independent map. Exploring the module maps together within the global map can be supported by Google Maps-based map navigation (discussed in the Section 5). The DNA map has a **modular structure** composed of eighteen interconnected functional modules, corresponding to ten DNA repair mechanisms, four phases of the cell cycle and four checkpoints, all interconnected, with multiple regulatory circuits (Figure 3A).

Due to the complexity of comprehensive networks, members of the same functional module can be very distantly located in the map. For example, a molecule that participates in several functional modules can have a geographical location in one module, but its location with respect to other modules can be far from optimal. To cope with this problem, we suggest creating a separate map for each module with an **individual module map layout**. The size of the module map is smaller compared to the global comprehensive map, and its contents are limited to the processes participating in the given module. This allows creating a layout where all entities of a single module map are closely located. This manipulation results in a different layout of the module, comparing it’s design at the comprehensive map, however it does not change the backbone structure of the module. An example of a module map with optimized individual layout is the Homologous Recombination (HR) from the DNA repair map, shown in Figure 3B. For modular map generation instructions see https://github.com/sysbio-curie/NaviCell.

Similarly, it is possible to generate **automatic layouts** for global, module maps or even for maps of an individual entity. The maps of an individual entity can represent ‘lifecycle’ reflecting reactions where the entity participates and the regulators of these reactions. This can be performed in Cytoscape software using the BiNoM plugin using the modularization and automatic layout functions (25). To facilitate the integration of separated signalling diagrams, there are at least two methods for map merging: (i) Merge Model plugin in CellDesigner (14) and (ii) BiNoM plugin on Cytoscape which allows to reorganize, dissect and merge disconnected CellDesigner pathway diagrams (25). For the map merging procedure in BiNoM see https://binom.curie.fr/docs/BiNoM_Manual_v2.pdf.

## Section 3: Data extraction and representation

### 3.1 Manual data mining

Retrieving knowledge and evidences from scientific papers, followed by a suitable graphical representation, requires a clear understanding of the biological processes and the experimental methodologies. Molecular interactions can be validated by various experimental methods. The most common and reliable experimental methods confirming different types of molecular interactions in the cell are summarized on Supplementary Table 3.

Depending on the purpose on the map, one can aim to represent biological processes with great precision including post-translational modifications, transport, complex association, degradation etc. Phenotype nodes on the signalling maps normally serve to indicate signalling readouts or cell statuses or biological processes in general. This type of nodes can also serve for schematic representation of statements when the exact molecular mechanism is still unknown. Some details on cell signalling might be skipped or, on the contrary, represented rigorously, depending on the purpose of the map being drawn, and the opinion of the map creator. It is important to address the challenge of **persistent homogeneity** on how to present the information on the map. This is especially important for visualisation and exploration of the map, because it will ensure a correct stepwise appearance of details at different zoom levels on the Google Maps-like map visualisation platform (discussed in the Section 5.2).

For efficient map construction, we suggest to perform a **systematic literature revision**. The hierarchical organization of the map, actually reflects a suggested approach for literature curation. First, we recommend to use **seminal review papers** in the field as well as the major **pathway databases** (Supplementary Table 1), in order to define the boundaries and general contents of the map. The information on canonical pathways retrieved from these sources, reflects the consensus view of the field and can serve as basis for drawing the backbone structure of the map. Thus, further details on molecular mechanisms can be compiled from the most recent studies and added to the map. The good practice is to comply with the requirement of supporting every interaction included in the map by at least **two independent investigations**.

In addition, the interpretation of scientific text and translation of written information into the correct and meaningful diagram is not always obvious. For consistency from text to diagram translation, we developed major rules for standardised interpretation of statements, see https://github.com/sysbio-curie/NaviCell. Figure 4 represents an example of scientific text translation into the PD diagram type.

**Figure 4.**
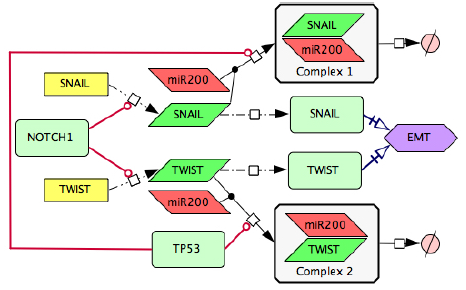
Transforming text to diagram: role of p53 and NOTCH in induction of epithelial to mesenchymal transition (EMT). The following statements were used for diagram construction: **1).** Control of EMT program is performed by SNAIL and TWIST, the major transcription factors that can induce the executors EMT program (51). These transcription factors are under the control of several upstream mechanisms. **2).** SNAIL and TWIST are inhibited by the p53 protein via a variety of microRNAs, including miR200 (52) **3).** miR20 binds to the mRNAs of SNAIL and TWIST and triggers their degradation, this way preventing the translation of mRNAs into the corresponding proteins (53) **4).** EMT program can be initiated due to excessive expression of NOTCH that directly activates transcription of SNAIL and TWIST (54).

### 3.1 Automatic data mining

The alternative approach to manual curation of scientific literature, is a wide range of automated or semi-automated text mining techniques (26). For example, the BioCreative initiative, an international community-wide effort for evaluating text mining and information extraction systems applied to the biological domain (http://www.biocreative.org). BioCreative gathers together multiple scientific teams and provides a list of text mining methods. In addition, it suggests common standards for text mining (27).

There are several automated text mining algorithms allowing to retrieve statements from scientific texts and to convert them into a meaningful graphical diagram. The advanced INDRA approach allows building computational models directly from natural language using automated assembly of network (28). The HiPub approach translates PubMed and PMC texts to networks of knowledge (29). The ANDSystem: an Associative Network Discovery System for automated literature mining, distinguishes and reconstructs in a form of networks several types of interactions as genetic, protein, metabolic, etc. (30). Finally, the Information Hyperlinked over Proteins (iHOP) system, uses genes and proteins as hyperlinks between sentences and abstracts, and converts the information from PubMed into one navigable resource for scientific literature investigation (31).

Among others, there is an approach of **formalized text-based intermediate language** as Biological Expression Language (BEL), that structures the statements in a way that can be mechanistically converted into a meaningful biological network (32). Similarly, BiNoM, a Cytoscape plugin, provides a function that converts sentences formulated in the “human” language, into a set of formal statements describing reactions that in turn can be automatically represented as graphical diagrams. The statements can be prepared in any simple text-based editor and imported into Cytoscape through BiNoM and converted into the CellDesigner diagram as presented in Supplementary Figure 1. The syntax of this BiNoM Reaction Format (BRF) language is depicted in (21)(33) and in the BiNoM manual https://binom.curie.fr/docs/BiNoM_Manual_v2.pdf.

## Section 4: Map annotation

### 4.1 Map entities annotation

Consistent integration of **standard common identifiers (ID)** for map entities ensures compatibility of maps with other tools and facilitates data integration into the maps. A common entity annotation form is needed for efficient map exploration by users and for map cross-curation by specialists in the corresponding fields. To address the challenge of systematic and user-friendly entity annotations, we developed the **NaviCell annotation format**. The entity annotation is normally performed by the map creator during the map construction in CellDesigner (Figure 3C).

We suggest to structure the annotation panel in four sections: “Identifiers”, “Maps_Modules”, “References” and “Confidence”. We provide an example of BRCA1 entity annotation from the DNA repair map in Figure 3C and Supplementary figure 2.

The **section “Identifiers”** includes standard identifiers (IDs) and provides links to the corresponding pages in HGNC, UniProt, Entrez, SBO, GeneCards and cross-references in REACTOME (34), KEGG (35), Wiki Pathways (36) etc. Metabolites and small compounds are annotated by corresponding IDs and linked to ChEBI (37), PubChem Compound (38) and KEGG Compound (39) databases.

The **section “Maps_Modules”** includes links to modules of the map where the entity participates. Referring to our example, the DNA repair map is part of Atlas of Cancer Signalling Network (ACSN) resource, and multiple entities from the DNA repair map can participate in various maps in the resource. There is a challenge to demonstrate this multi-functionality of an entity and there is a need to provide links to all maps and modules of ACSN where an entity participates. To achieve this, we introduced a tagging system, included into the entity annotation. The tags with map and module names where the entity participates are systematically included into the annotation during the map construction in CellDesigner.

The **section “References”** provides links to the corresponding publications from which the evidences on the entity or its reactions were collected. In addition, map creator’s notes can be added in this section, in order to emphasize or to highlight important information related to the entity or its reactions. Each entity annotation is represented as a web page (Supplementary Figure 2) automatically generated when the NaviCell map is generated from the CellDesigner file (described in the Section 5.1). The extended description of annotation formats for each type of entities in the map is provided in the documentation, see https://github.com/sysbio-curie/NaviCell.

### 4.2 Confidence scores

Despite that multiple network and pathway resources are believed to summarise the current knowledge on biological processes, there is always a remaining open question: to what extent the retrieved statements from the scientific literature and depicted in the diagrams, are indeed reflecting the biological reality? There is a permanent problem that map creators need to address: what processes should be included in the map diagrams? The challenge is how to systematically evaluate the experimental evidence from scientific publications and assign a confidence score. There is no consensus about the approach to evaluate the “truth” or the “degree of confidence” regarding to molecular mechanisms reported in the literature.

Here, we provide several examples of confidence scores developed in well-established signalling networks resources. The scores in the BioGRID resource (40) (https://thebiogrid.org) are calculated taking into account if it is an original publication describing the interaction following the principle of CompPASS score system that uses unbiased metrics to assign confidence measurements to interactions from parallel nonreciprocal proteomic data sets (41). These scores are ranged from 0 to 1, with 1 being the most confident. The SIGNOR database (42) (SIGNOR http://signor.uniroma2.it) uses functional relevance of interactions provided in the form of a “reliability score”. This score is defined by the probability that two partners in the interaction are cited together in the same paper. This co-citing probability is ranged from 0 to 1. In the RECON database (43) (https://vmh.uni.lu) each reaction in the network is associated with a confidence score ranging from 0 to 4. This score is based on the type of evidence identifying the reaction, assigning values for modelling (score 1), sequence (score 2), physiological (score 2), genetic (score 3) and for biochemical (score 4) studies. If multiple evidence types exist for the same interaction, a cumulative confidence score is assigned.

To evaluate the reliability of depicted molecular interactions in the ACSN resource, there are **two confidence scores** for protein complexes and reactions. The scores are automatically calculated during the conversion of the CellDesigner map to the NaviCell format and provided in the annotation’s **section “Confidence”,** in a form of a five-star diagram (Figure 3C and Supplementary Figure 2). Both scores are integers, ranging from 0 (undefined confidence) to 5 (high confidence).

The reference score, marked by “REF” indicates both, the number and the “weight” associated to publications found in the annotation of a given reaction, with weight equal 1 point for an original publication and 3 points for a review article. For example, a reaction annotated by two reviews will obtain the highest reference score REF = 5. The functional proximity score, marked by “FUNC” is computed based on the external network of protein-protein interactions (PPI), Human Protein Reference Database (HPRD) (44). The score reflects an average distance in the PPI graph between all proteins participating in the reaction as reactants, products or regulators. Therefore, if all the proteins participating in a reaction interact directly (functional distance 1) in the PPI network, then this reaction obtains the highest score (FUNC=5). If in the reaction where phosphorylation of protein A is catalysed by protein B, and A and B are separated in PPI graph by 2 (or 3, or 4) edges, then the reaction obtains the functional proximity score FUNC=4 (or FUNC=3, or FUNC=2 respectively). If two reaction participants are not connected in the PPI network, then FUNC=0. The functional proximity is computed using BiNoM, Cytoscape plugin (25).

## Section 5: Map generation and navigation in NaviCell

### 5.1 Generation of NaviCell map using NaviCell factory

CellDesigner maps, as the DNA repair map, annotated in the NaviCell format or not, can be converted into a NaviCell web-based front-end, which represents a set of html pages with embedded JavaScript code. It can be launched in a web browser locally or located on a web-server. The NaviCell factory is currently embedded in the BiNoM Cytoscape plugin but will be soon available as a stand-alone command line package. The detailed guide to use the NaviCell factory is provided at https://github.com/sysbio-curie/NaviCell.

### 5.2 Common exchange formats

The generated maps in CellDesigner and exposed in the NaviCell format, can also be provided in **common exchange formats,** as BioPAX and PNG formats. In addition, the modular composition of the maps can be provided in the form of GMT files. The description of map preparation in various formats using the BiNoM Cytoscape plugin is available in https://binom.curie.fr/docs/BiNoM_Manual_v2.pdf.

### 5.3 Map navigation using NaviCell platform

The problem of navigation and exploration of modern digital interactive geographical maps is successfully addressed in Google Maps or similar systems. Large molecular interaction maps are comparable in size and complexity to geographical maps, therefore the navigability challenge can be addressed in a similar manner.

Comprehensive signalling maps, as the DNA repair map, contain a large number of nodes and edges, leading to a complex map navigation. To solve this problem, maps can be represented as clickable web pages with a user-friendly interface. We developed the NaviCell web-based environment, empowered by Google Maps engine for visualization and navigation of molecular maps (18). Scrolling, zooming, markers, callout windows as well as a zoom bar, are adopted from the Google Maps interface (Figure 3 and Supplementary Figure 3). The map in NaviCell is interactive and all map components are clickable. To find an entity of interest or querying for a single or multiple molecules is possible using the search window. Alternatively, the entity can be found in the selection panel or directly on the map.

### 5.4 Map navigation modes

The NaviCell platform is convenient for several map navigation modes. The so called **horizontal navigation** is facilitated by the hierarchical modular structure of the maps, which is successfully exploited by NaviCell system. The modular structure of maps allows to transfer from the global map to the module maps. Thus, using our example of the DNA repair map, the involvement of BRCA1 protein across processes on the map can be appreciated (Figure 3A). However, in order to understand the details of the biochemical reactions where BRCA1 plays a role in a particular process as Homologous recombination, the modular map is more suitable (Figure 3B).

The so called **vertical navigation** is facilitated by the semantic zooming feature of maps prepared in the NaviCell format. **Semantic zooming** (see Table 1 for terms and definitions) simplifies the navigation through the large maps of molecular interactions, showing a readable amount of details at each zoom level. This gradual detail appearance permits to explore the map contents from the top-level toward a detailed view. To prepare maps for this type of navigation, a map pruning is performed, eliminating hence non-essential information for each zoom level. Typically, it is recommended to generate three to four zoom levels, although the number of zoom levels is unlimited in NaviCell (Figure 5).

**Figure 5.**
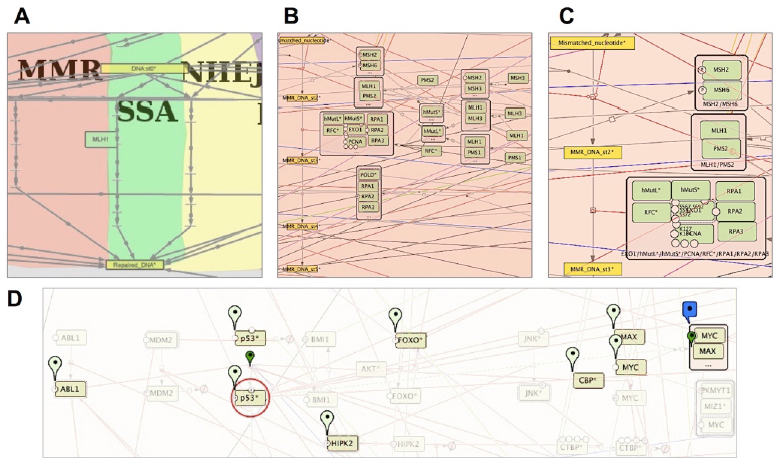
Semantic zooming and entity visualization on DNA repair map. (A)Canonical pathways view, (B) Hide-details view, (C) Detailed view, (D) Highlighting p53 neighbours.

In our example of the DNA repair map, the top-level zoom displays the modules of the map, depicted with a coloured background shapes with the backbone structure of the pathways (Figure 5A). In the next zoom level, canonical cell signalling pathways are shown. These pathways are defined by intersecting the content of the map with the corresponding pathways in other databases (Figure 5B), at the last zoom all map elements are present (Figure 5C). For detailed instructions on map zoom levels creation see https://github.com/sysbio-curie/NaviCell.

An additional way for map exploration is **highlighting individual entities** of the map and their **neighbours**. We developed a NaviCell function for selecting species of interest and its neighbours. This function allows step-wise enlarging of the neighbourhood coverage to understand the signalling propagation on the map, as shown for p53* molecule on the DNA repair map, in Figure 5D.

## Section 6: Map maintenance and curation

Biological knowledge about the majority of signalling pathways is not yet solid and grows continuously. Due to this, one of the major problems of signalling maps is their fast obsolescence. To address this challenge, permanent maintenance and map updating is needed. The community of users is the most reliable and trustable contributor to map maintenance, because specialists can support and update maps from their own research area. To enable such a community-based effort, efficient curation tools should be created. To our knowledge, there is only one community curation tool for comprehensive maps, the Payoa plugin of CellDesigner (45).

We suggest carrying out map curation in the context of NaviCell environment via a blog. The process of map curation and maintenance in NaviCell involves map managers that check the posts of the maps in the blog and latest scientific literature, and update the maps accordingly. An automated procedure supports the map updating and archives older versions of posts, including comments, providing thus traceability of every map changes and all blog discussions (18).

## Section 7: Visualisation of omics data in the context of signalling network maps

To make data visualisation a straightforward and easy task, we developed a **built-in toolbox for visualisation and analysis of high-throughput data** in the context of comprehensive signalling networks. The integrated NaviCell web-based toolbox allows importing and visualising heterogeneous omics data on top of the maps, and to perform simple functional data analysis. standard common identifiers (IDs) in the annotation of proteins, genes (HUGO) and metabolites (CHEBI) allows the NaviCell data visualisation functionality. NaviCell also computes aggregated values for sample groups and protein families. For visualization, the tool contains standard heatmaps, barplots and glyphs, as well as the map staining technique for displaying large-scale trends in the numerical values on top of the map. The combination of these flexible features provides an opportunity to adjust the modes of visualisation to the type of data, and to acquire the most meaningful picture (19). Extended documentation, a tutorial, a live example and a guide for data integration using NaviCell are provided at https://navicell.curie.fr/pages/nav_web_service.html.

To illustrate data visualization, the EMT regulation map from NaviCell collection (9) https://navicell.curie.fr/pages/signalling_network_emt_regulation_description.html was used to analyse omics data from ovarian cancer patients (46). The data files can be downloaded on https://navicell.curie.fr/pages/nav_web_service.html.

Expression and mutational profiles of proliferative and mesenchymal ovarian cancer classes were compared. Expression data from the two groups of patients was visualised on the EMT regulation map using the map staining technique in NaviCell, showing average expression of each gene across the samples. The mutation profiles for the same disease classes were visualised using glyphs. There is a clear difference in the expression pattern between the two classes, indicating a major activation of epithelial to mesenchymal transition (EMT) regulators in mesenchymal class (Figure 6B), characterised by invasive clinical outcome, opposite to the proliferative class (Figure 6A). A closer look at the EMT inducers ZEB1&2, SNAIL, CTNBB1, shows that they are largely mutated and demonstrate a low expression pattern in the proliferative, clinically, less invasive class (Figure 6C), in opposite to the invasive mesenchymal class (Figure 6D). The observation is consistent with the notion that in order to induce ETM and invasion, the cancer cell needs to be in low-proliferative status (9). Similar molecular portraits of cancer can be automatically generated using NaviCom, that connects cBioPortal and NaviCell allowing visualisation of different high-throughput data types simultaneously on a network map in one click (22).

**Figure 6.**
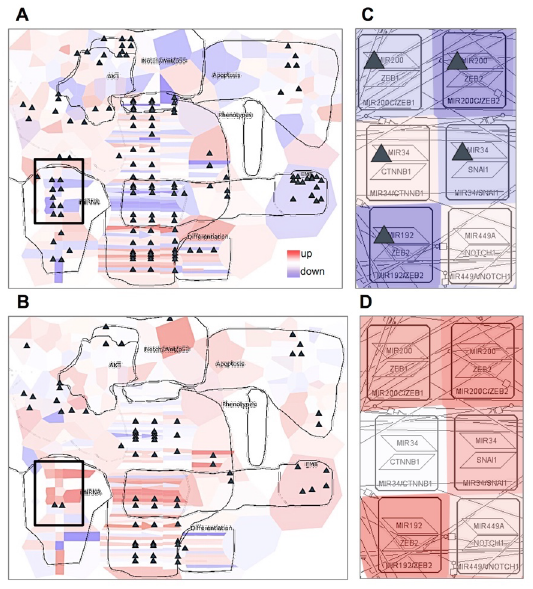
Visualization of cancer high-throughput data in the context of DNA repair map. Visualisation of gene expression from ovarian cancer samples in a form of map staining and mutation profile in a form of glyph (triangle). (A) Proliferative and (B) Mesenchymal classes of ovarian cancer. Zoom in on EMT regulators in (C) Proliferative and(E)Mesenchymal classes of ovarian cancer. Proliferative group, n=87, Mesenchymal group, n=96.

Maps generated using the procedure described above can be applied in different studies, not exclusively in NaviCell environment. For instance, the EMT regulation map (Figure 7A) was used to retrieve a minimal mechanistic model explaining the control of the EMT program. With this aim, hub players in each functional module of the map were identified (Figure 7B) and network complexity reduction was performed using path analysis function in BiNoM Cytoscape plugin (25) up to core regulators of EMT, apoptosis and proliferation, that were preserved through all levels of reduction (Figure 7C). The minimal model was used to predict genetic interactions leading to invasive phenotype in colon cancer (9).

**Figure 7.**
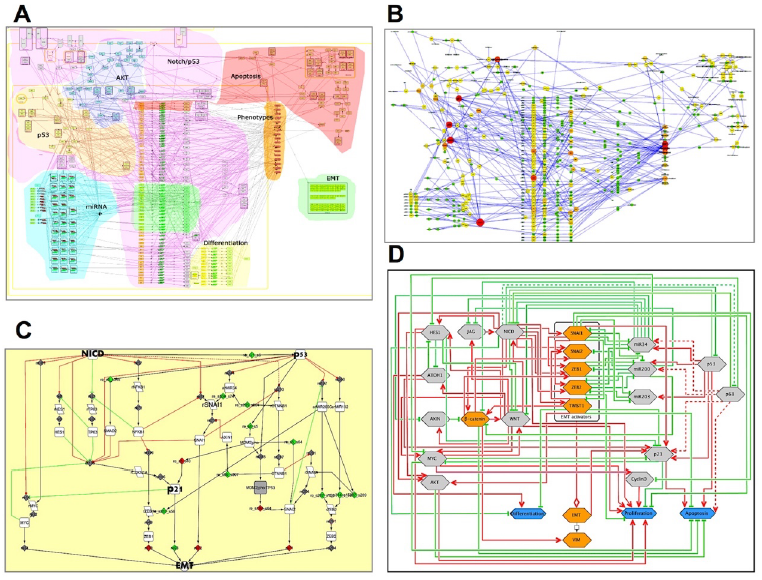
Retrieval of mechanistic model of EMT program control using EMT regulation map. (A) Comprehensive signalling map of EMT regulation, (B) Hub player in functional modules of EMT map, (C) Structural analysis and reduction of EMT map complexity; (D) Scheme representing major mechanisms controlling EMT program.

## Conclusions and perspectives

The knowledge about biological mechanisms is rapidly growing, demanding organisation and systematic representation of the information to create a global picture on cell molecular mechanisms (47). The current solution is to visualise cell signalling in a form of a diagram or a molecular map. The maps can vary in their size, complexity, contents and details’ description. However, the majority of the map construction endeavours to address very similar challenges. Among others, the questions on how to improve the standardised representation of biological processes, provide intuitive map navigation tools, optimise maps update with the latest literature, are still not fully solved.

In this manuscript we described the methodology developed following the long-standing experience with comprehensive generation and manipulation of maps. Using this approach we created and currently maintain the Atlas of Cancer Signalling Network (ACSN) resource (20) and a collection of maps created in CellDesigner available at https://navicell.curie.fr/pages/maps.html. We suggested a workflow for construction and annotation of signalling maps in CellDesigner, preparing hierarchical modular structure of maps and also generation of different map view levels, to allow semantic zooming-based exploration of maps in NaviCell. We introduced NaviCell, an environment for navigating large-scale maps of molecular interactions created in CellDesigner. NaviCell allows showing the content of the map in a convenient way, at several scales of complexity. It provides an opportunity to comment the map contents, thus facilitating its curation and maintenance. Finally, we showed how omics data can be visualised and interpreted in the context of the map.

Among many future challenges for the signalling network community, is the integration of similar efforts. Actually, the open-flow model of knowledge sharing and integration was suggested in the past by Kitano and colleagues (48). Nowadays, there are several examples of such efforts as Wiki Pathways (49) and Pathway Common (50). In addition, there is an emerging community activity, the Disease Maps Project, where mechanisms involved in various human diseases are brought together, allowing disease comorbidity and drug repositioning studies (http://disease-maps.org). Finally, the generation of comprehensive platforms for tools, data, and knowledge sharing in systems biology and biomedical research, similar to GARUDA initiative (http://www.garuda-alliance.org), will facilitate the compatibility between tools and resources.

The map construction and annotation procedures that we expose in this manuscript aim to integrate community efforts, which are in line with FAIR principles (23). We are convinced that the application of standardised map construction procedures, providing access to the maps’ content via web-based platforms, will increase the reuse of the existing maps, improving management and updating, and will allow efficient generation of new high-quality signalling maps. Actually, the idea of exchangeable, re-usable maps or map modules lies in the basis of the Disease Community resource creation, where common mechanisms implicated in various diseases will be shared between disease-specific maps. Finally, standardly created maps provided in common formats will ensure their compatibility with various systems biology tools, facilitating their applications for data analysis and modelling.

## Acknowledgements

We thank L. Cristobal Monraz Gomez for critical reading of the manuscript. This work has been supported by the COLOSYS grant ANR-15-CMED-0001-04, provided by the Agence Nationale de la Recherche under the frame of ERACoSysMed-1, the ERA-Net for Systems Medicine in clinical research and medical practice and by INSERM Plan Cancer N° BIO2014-08 COMET grant under ITMO Cancer BioSys program. This work received support from MASTODON program by CNRS (project APLIGOOGLE).

## Supplementary materials

**Supplementary Table 1:**
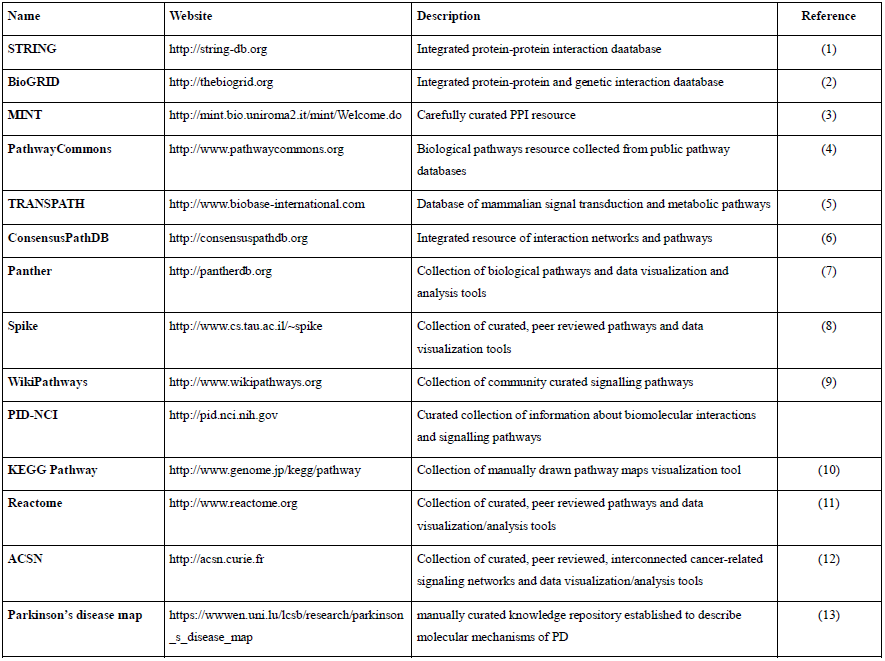
Pathways databases and network resources.

**Supplementary Table 2:** Biological diagram editors, map navigation tools and high-throughput data visualization support.

| Biological diagram editors |  |  |  |
| --- | --- | --- | --- |
| Name | Website | Description | Reference |
| CellDesigner | http://www.celldesigner.org | Structured diagram editor for drawing gene-regulatory and biochemical networks | (14) |
| SBGN-ED | http://vanted.ipk-gatersleben.de/addons/sbgn-ed | VANTED add-on for create and edit three types of SBGN maps | (15) |
| CellPublisher | http://cellpublisher.gobics.de | KEGG database-associated tool for data visualization and analysis in the context of pathway maps | (16) |
| Cytoscape / BiNoM | http://www.cytoscape.org<br>http://binom.curie.fr | Software platform for manipulation of biological networks represented in standard systems biology formats | (17)<br>(18) |
| Payao | http://payao.oist.jp:8080/payaologue/index.html | Network curation tool for simultaneous map commenting using tag system | (19) |
| NaviCell | http://navicell.curie.fr | Web-based tool for heterogeneous data visualization and analysis in the context of signaling networks | (20)(21) |
| yEd graph editor | http://www.yworks.com/en/products/yfiles/yed/ | Application for generate high-quality diagrams construction | http://link.springer.com/chapter/10.1007/978-3-642-18638-7_8 |
| VisANT | http://visant.bu.edu/ | Tool for visual analyses of metabolic networks in cells and ecosystems | (22) |
| Pathway Map Creator | http://lifesciences.thomsonreuters.com/m/pdf/PathwayMapCreator-cfs-en.pdf | Tool for editing and analysis of canonical pathways maps |  |
| Tools for visualisation of high-throughput data in the context of signalling networks |  |  |  |
| Name | Website | Description | Reference |
| ReactomeFiViz | http://wiki.reactome.org/index.php/Reactome_FI_Cytoscape_Plugin | Cytoscape plugin for data integration into signaling networks | (23) |
| iPath | http://pathways.embl.de | Web-based tool for data visualization in the context of pathway maps | (24) |
| Medusa | http://coot.embl.de/medusa | Tool for data visualization in the context of signaling network and network clustering | (25) |
| NaviCell | http://navicell.curie.fr | Web-based tool for heterogeneous data visualization and analysis in the context of signaling networks | (20)(21) |
| NaviCom | http://navicom.curie.fr | Web-based platform for generating interactive network based molecular portraits using high-throughput datasets. | (26) |
| MINERVA | http://r3lab.uni.lu/web/minerva-website | Webserver for visualization, exploration and management of molecular networks encoded in SBGN-compliant format | (27) |
| KEGG Mapper | http://www.kegg.jp/kegg/mapper | KEGG database-associated tool for data visualization and analysis in the context of pathway maps | (10) |

**Supplementary Table 3:** Method for studying molecular and genetic interactions.

| Interaction | Method | Reference |
| --- | --- | --- |
| **Ligand-receptor interactions** | Resonance energy transfer (FRET and BRET) | (28)(29) |
|  | Flow Cytometric Analysis | (30) |
| **Direct protein-protein interactions** | Co-immunoprecipitation (CoIP), | (31)(32) |
|  | NMR, X-ray crystallography, | (33) |
|  | GST-pull down assay | (34) |
|  | Tandem affinity purification | (31)(29) |
|  | Far Western blotting | (35) |
|  | Phage display | (32) |
|  | Mass-spectrometry | (36) |
|  | Two hybrid assays (yeast and mammalia) | (31) |
|  | Functional mutational analysis | (37) |
| **Direct protein-DNA interaction (transcription regulation and co-regulation)** | Chromatin immunoprecipitation (ChIP) | (38)(39) |
|  | DNA footprinting, | (38) |
|  | Electrophoretic mobility shift end supershift assays (EMSA) | (38) |
|  | Computational prediction of transcription factors binding sites | (40) |
| **MicroRNA binding** | Direct miRNA binding assay, | (41) |
|  | 3'UTR reporter assay, | (41) |
|  | Computational miRNA target prediction | (41) |
| **Regulation of expression (mRNA and protein level)** | Reverse transcription polymerase chain reaction (RT-PCR) | (42) |
|  | Reporter assays | (43) |
|  | RNase protection assay | (44) |
|  | Nothern blot | (42) |
|  | Western blot | (45) |
|  | Fluorescence-activated cell sorting (FACS) | (46) |
| **Genetic interactions** | Genetic knock-out, knock-down, knock in or overexpression of effector molecules | (47)(48) |
|  | Synthetic interaction detection assays | (32)(49) |

**Supplementary Figure 1.**
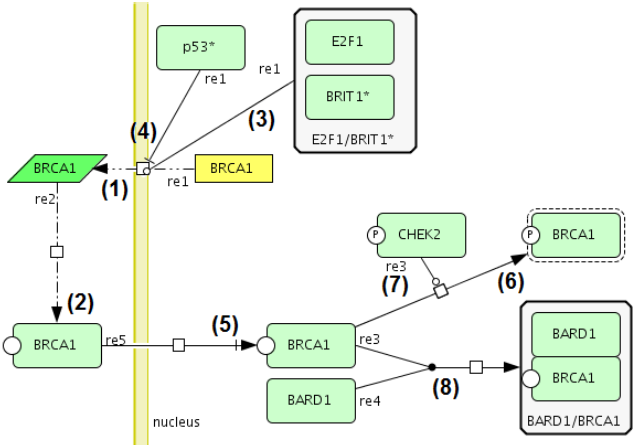
Automatic text to diagram translation using BiNoM Reaction Format (BRF) language.

Representation of biochemical reactions from the following statements. Numbers correspond to the reactions in the diagram: *« BRCA1 transcription (1) and translation (2) is positively regulated by E2F1/BRIT1* complex (3) and inhibited by p53 (4). BRCA1 protein is transported into nucleus (5), where CHEK2 kinase activates it by specific phosphorylation (6) and (7). Additionally, BRCA1 forms a complex with BARD1 (8) and BRCA1 association with BARD1 is essential for the E3 ligase activity of BRCA1».* References correspondence: reactions 1,2,4 (50); reaction 3 (51); reaction 5 (52); reactions 6,7 (53); reaction 8 (54).

The set of corresponding statements in BiNoM Reaction Format (BRF) language were generated from the text and translated into the diagram:

BRCA1@cytoplasm -/> BRCA1@nucleus

BRCA1@nucleus+BARD1@nucleus-:> BARD1:BRCA1@nucleus

BRCA1-CHEK2|pho@nucleus -> BRCA1|pho|active@nucleus

rBRCA1@cytoplasm -.> BRCA1@cytoplasm

gBRCA1@nucleus-|p53*@nucleus-BRIT1*:E2F1@nucleus-..>rBRCA1@cytoplasm

**Supplementary Figure 2.**
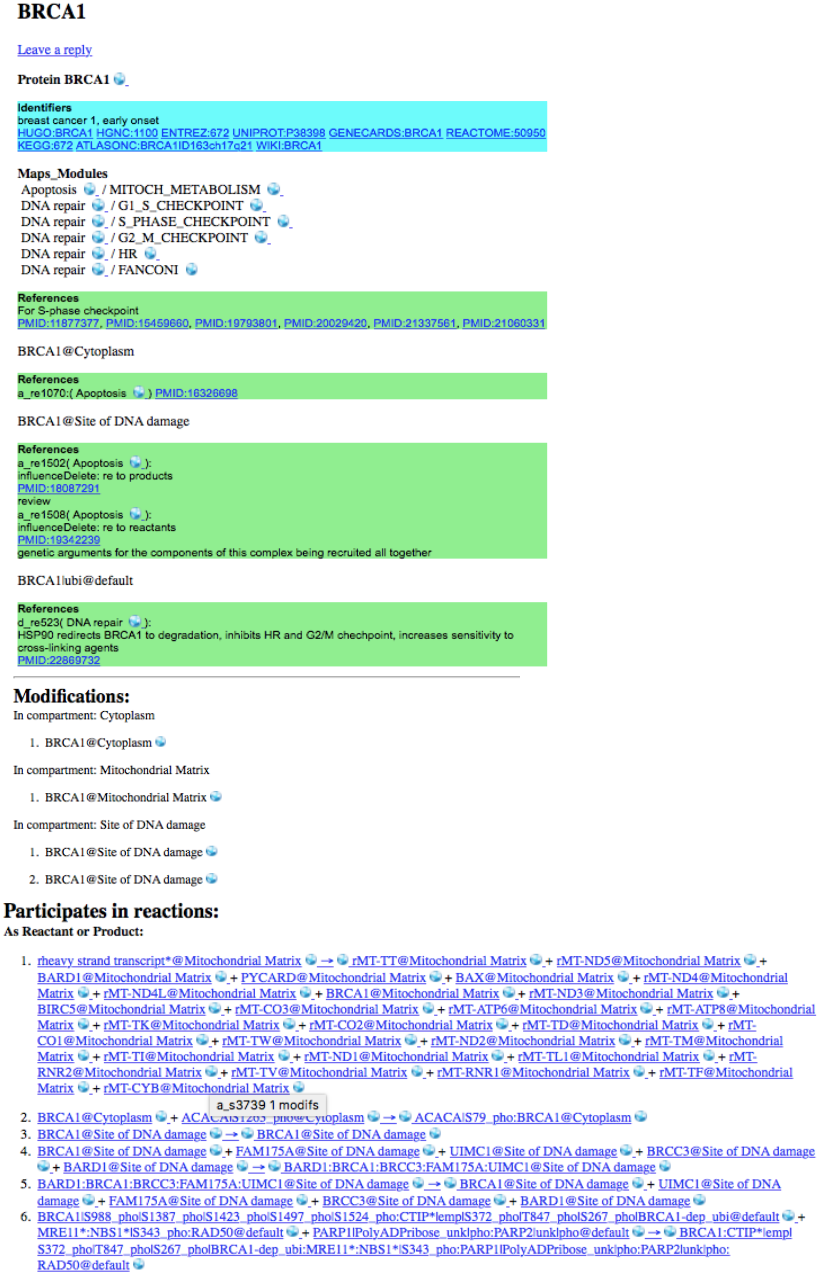
Annotation page in the ACSN blog for BRCA1 protein.

## Access, documentation and tutorials

**Maps construction:**

**CellDesigner introduction and tutorial**

http://celldesigner.org/documents.html

**SBGN**

http://www.sbgn.org/Main_Page

**BiNoM manual**

https://binom.curie.fr/docs/BiNoM_Manual_v2.pdf

**Preparing modular map in NaviCell format and converting to NaviCell web-based environment:**

https://github.com/sysbio-curie/NaviCell

**Data visualization and analysis:**

**NaviCell Web Service introduction, tutorial and case studies**

https://navicell.curie.fr/pages/nav_web_service.html

**NaviCell Web Service guide**

https://navicell.curie.fr/doc/ws/NaviCellWebServiceGuide.pdf

**Interactive demo on data visualization using NaviCell**

https://navicell.curie.fr/navicell/maps/cellcycle/master/index.php?demo=on

**NaviCom guide**

https://navicom.curie.fr/tutorial.pdf

**Maps access:**

**ACSN introduction, tutorial and case studies**

https://acsn.curie.fr/documentation.html

**NaviCell maps collection**

https://navicell.curie.fr/pages/maps.html

